# Ecophylogenetic patterns of rhizosphere bacterial community assembly in *Pisum* spp. (Fabaceae, Fabeae) reveal strong plant-mediated ecological filtering

**DOI:** 10.1101/2025.09.30.679227

**Authors:** Victor Angot, Vincent Pailler, Abdelhamid Kebieche, Elodie Belmonte, Virginie Bourion, Yanis Bouchenak-Khelladi

**Affiliations:** Université Bourgogne Europe, Institut Agro, INRAE, UMR Agroécologie, 21000 Dijon, France; INRAE, UMR GDEC, Gentyane, Clermont-Ferrand, France; Institut Agro, Université d’Angers, INRAE, IRHS, SFR 4207 QUASAV, Angers, France

**Keywords:** plant-microorganisms interactions, *Pisum* spp, plant development, ecophylogenetic, ecological filtering

## Abstract

Plant-microorganisms interactions are among the oldest biotic relationships and play a fundamental role in shaping biological systems. These associations involve several and diverse species, each evolving on different timescales. Understanding these interactions requires approaches that integrate both ecological community dynamics and evolutionary processes, which drive the adaptation of plants and microorganisms. We investigated bacterial community assembly dynamics associated with four diverse *Pisum* spp. accessions grown in greenhouse conditions on soil. Bacterial DNA was extracted from bulk soil, rhizosphere, rhizoplane, and endosphere microhabitats across three plant growth stages, followed by full-length 16S rRNA gene sequencing. Bacterial communities varied in diversity, composition and structure across microhabitats and growth stages. Ecophylogenetic analyses, that is integrating community ecology dynamics into a phylogenetic framework, indicated strong host-filtering, with community assembly across space and time being structured by phylogenetic constraints. This illustrates the role of the plant in creating and shaping distinct ecological niches, where selective recruitment favors specific and closely related lineages. Our findings suggest that an ecophylogenetic approach provides valuable insights into plant-microbiota dynamics by integrating ecological and evolutionary processes, thereby offering a powerful perspective to investigate the co-adaptation of plants and their associated microbiota.

## Introduction

Biotic interactions are responsible for major evolutionary changes, such as increased genetic diversity within populations or triggering adaptive changes leading to ecological speciation, across multiple spatial and temporal scales. These complex interactions, that can involve two or thousands of species and occur within or across trophic levels, contribute to crucial ecological patterns that structure biological systems (Schemske *et al.*, 2009; Spicer & Woods, 2022). One of the most intricate and complex biotic interaction is the association between plants and microorganisms (Lyu *et al.*, 2021), that is among the oldest relationship that may have originated around 460-475 Mya (Brundrett, 2002), and represents an ideal model for studying host-microbiota association dynamics.

Legumes (Fabaceae) is a highly diverse family of angiosperms, many of which form symbiotic associations with nitrogen-fixing bacteria and play an important role in fixing atmospheric nitrogen. Globally, grain legumes (tribe Fabeae) were estimated to fix a total of 35.5 million tons of nitrogen every year (Herridge et al., 2022). Among these, *Pisum* spp. include one of the first domesticated plant species (Zohary & Hopf, 1973). *Pisum* is divided into two species: *P. fulvum* and *P. sativum* which is subdivided in 3 sub-species: *P. sativum* subsp. *sativum, P. sativum* subsp. *elatius* and *P. sativum* subsp. *abyssinicum* (Smýkal et al., 2011). While the symbiotic relationship of *Pisum* spp. with *Rhizobia* is well-documented (Bourion *et al.*, 2018; Hawkins & Oresnik, 2022), the complex interactions with non-symbiotic microorganisms remain largely unknown, whereas the rhizosphere microbiota play a crucial role in shaping *Rhizobia*-legumes interactions (Han *et al.*, 2020). The non-symbiotic bacteria could also improve the effectiveness of *Rhizobia*-inoculants by enhancing their ability to colonize, survive, and persist in the soil, as well as their capacity to interact with other microorganisms (O’Callaghan et al., 2022; Kong & Liu, 2022). Therefore, it is critical to understand the assembly rules of the whole bacterial community, beyond *Rhizobium* symbiosis. Plant-microorganisms interactions are governed by a complex ‘chemical dialogue’ through the production of exudates by the plant (primary and secondary metabolites) (Duc *et al.*, 2022). Some exudates, such as organic acids (Rekha *et al.*, 2018), amino-acids (Feng *et al.*, 2018) and glucose (Zhang *et al.*, 2015), have chemo-attractive properties that facilitate bacterial colonization (Chaparro *et al.*, 2013). Other compounds, such as antimicrobial peptides, have antibiotic properties that prevent the colonization of pathogenic microorganisms (Pacheco-Cano *et al.*, 2018). In addition, root architecture and root hairs structure further modulate the assembly of the rhizosphere microbial communities (Kawasaki *et al.*, 2016; Robertson-Albertyn *et al.*, 2017) by altering the physical microhabitat properties. Therefore, through these chemical and physical modifications, plants can strongly influence the composition of rhizosphere microbial communities, exceeding the influence of the original soil properties (Tkacz *et al.*, 2020). In return, these shaped communities can enhance plant fitness through nitrogen fixation, nutrient acquisition, and increased resistance to biotic and abiotic stresses (Bhattacharyya & Jha, 2012; Peix *et al.*, 2015; de la Fuente Cantó *et al.*, 2020; Yang *et al.*, 2021).

The assembly dynamics of microbial communities are shaped by multiple factors, requiring multifactorial analyses that account for both spatial and temporal dimensions. Different plant compartments (referred hereafter as plant-associated microhabitats), such as the rhizosphere, rhizoplane and endosphere contain distinct microbial communities (Cregger et al., 2018; Gupta et al., 2021). Therefore, the identification of microbial associations and enrichment or impoverishment dynamics through an ecological gradient (i.e., from the rhizosphere to the endosphere) is crucial in order to characterize the host-mediated ecological filters associated with different plant-associated microhabitats (Zhang et al., 2017; Iven et al., 2022). Studies have shown that plant-associated microbial communities are specific to their host species and can shift through plant growth (Chaparro *et al.*, 2014; Edwards *et al.*, 2015; Shade *et al.*, 2017), suggesting both spatial and temporal filtering. While it is demonstrated that assembly of soil microbial communities is determined by biotic interactions such as mutualism (Goberna et al., 2014; Romdhane et al., 2022), facilitation or competition (Faust & Raes, 2012), the association and assembly dynamics throughout plant-associated microhabitats and plant growth remain largely unknown (Moroenyane *et al.*, 2021). To decipher these spatio-temporal dynamics, an ecophylogenetic framework is particularly effective because it computes ecological dynamics while considering phylogenetic relatedness to infer underlying biotic interactions. For instance, insights from human microbiomes indicate that microbial recruitment can follow an underdispersion assembly pattern (Darcy *et al.*, 2020) (i.e. taxa are more likely to be recruited if a phylogenetically close relative was previously established). Whether similar processes govern the mode and tempo of microbial recruitment in plant-associated microhabitats remains largely speculative. The use of ecophylogenetic frameworks is therefore needed to reveal association patterns with host plants (Wang & Sugiyama, 2020). The assembly of these community may be attributed to phylogenetic niche conservatism (Losos, 2008; Pyron et al., 2015), when closely related species inhabit similar ecological niches. However, instances of competitive exclusion have been shown, such as in floral nectar (Peay *et al.*, 2012), and the balance between these processes may be influenced by resource availability overtime (Pekkonen et al., 2013). Similarly, the intensity of plant rhizodeposition and its ability to shape ecological niches for bacteria could affect the pattern of phylogenetic association. The specialization of microbiota on the plant’s physiological needs and on its microhabitats suggest that plant-microbiota interactions prioritize specific functions over particular microbial taxa (Castellano-Hinojosa & Strauss, 2021). Since phylogenetic relatedness often predicts functional similarity (Burns & Strauss, 2011; Martiny *et al.*, 2015; Anacker & Strauss, 2016), community assembly is expected to be phylogenetically structured (Aguirre de Cárcer, 2019). Therefore, an ecophylogenetic approach can help identify whether plant-mediated ecological filtering shapes microbial assembly across spatial and temporal scales.

To explicitly link spatial and temporal host-mediated ecological filters with phylogenetic community assembly, we conducted a greenhouse experiment with four *Pisum spp.* accessions to study the composition and structure of plant-associated belowground bacterial communities at different growth stages using both ecological and phylogenetic approaches. Our hypotheses were (i) that bacterial community diversity, composition and phylogenetic diversity differ across the plant-associated microhabitat and reflect a gradient of host-filtering, (ii) that bacterial community composition and phylogenetic structure shift across plant growth stages, reflecting dynamic changes in host-filtering, and (iii) that host-filtering results in non-random phylogenetic assembly patterns.

## Material and Methods

### Experimental design and data collection

Four *Pisum* accessions from the Pea Genetic Resource Centre at INRAE Dijon were selected: (i) one *Pisum fulvum* (IPIP10023) and (ii) one *Pisum sativum* subsp*. elatius* (IPI200733) as two wild species accessions, one *Pisum sativum* subsp*. abyssinicum* ‘ATAR’ (IPI200596) and one *Pisum sativum* subsp*. sativum* ‘CAMEOR’ (IPI200293) as two accessions representative of domestication or selection events. The seeds were stored in a cold room prior to sowing. Seeds from *Pisum fulvum* and *Pisum sativum* subsp*. elatius* were scarified using a scalpel to enable germination. Soil was collected at Auxonne, Côte d’Or, France (47.185N, 5.403E) from an organic farm, air-dried (Han *et al.*, 2026), and sieved to 4 mm. Soil nitrate, ammonium, and total nitrogen, as well as water content and bulk density, were assessed at the laboratory “SADEF Agronomy & Environment” (Appendix 1). The experiment was conducted as a completely randomized design and was carried out in a climate-controlled glasshouse located at INRAE Dijon from February to May 2023. The greenhouse was equipped with a temperature control system that maintained a temperature of 20 °C during the day and 15 °C at night, and a 16 hours photoperiod. Seeds were sown into sterilized 2 L pots filled with sieved soil. Two seeds were sown in each pot to ensure the successful germination of at least one. Pots were top-watered with osmotic water with an automatic dripper system until germination. Afterwards, pots were watered daily with 100 mL of a nutritive solution (MgSO4 + 7H2O 1 mL/L, CaCl2 0.63 mL/L, K2SO4 1.39 mL/L, NaCl 0.1 mL/L, Oligo 1 mL/L, and iron versenate 1 mL/L; pH 6.5). Plants were harvested at three successive growth stages: the first stage (S1) was the vegetative stage (5 to 6 leaf nodes, 16-24 days), the second stage (S2) was the beginning of flowering (appearance of the first flower, 31-56 days); and the third stage (S3) was the beginning of seed filling (the first pod containing internal seeds at least 8mm in length, 36-64 days). The aim was to have nine plants for each growth stage and for each accession (nine replicates x three growth stages x four accessions) resulting in a total of 108 plants, with 27 plants per accession. However, not all seeds germinated, with only 25/27 for *Pisum sativum* subsp*. abyssinicum,* 21/27 *for P. sativum* subsp*. sativum, 15/27 for P. fulvum and 15/27 for P. sativum* subsp*. elatius*, which led to a total of 76 plants. Eight pots were left unsown to provide bulk soil samples without any association with *Pisum*, serving as controls. After retrieving the plant, the roots were transferred to a sterile centrifuge tube with 35 mL of 1X PBS and kept at 4 °C. Roots were initially sonicated for one minute and 30 seconds to collect rhizosphere microorganisms (R), then transferred to a new tube containing 35 mL of 1X PBS and subjected to a second round of sonication for 10 minutes to isolate rhizoplane microorganisms (Rp). The R and Rp tubes were then centrifuged at 5000 x g for 10 minutes, and 250 mg of the resulting pellet was sampled for DNA extraction. Additionally, 400 mg of the central root pivot (the first 3 cm below seed insertion) were sampled and grinded to retrieve the endosphere (E) microbial communities. At the start of the experiment, and at each stage, bulk soil samples were collected as controls and analyzed using the same procedure as for R and Rp. Some plant nodules were also sampled, however, due to the low number of plants that formed nodules, the results concerning this microhabitat could not be included in this study. In total, we collected 260 samples that encompassed all plant-associated microhabitats at three growth stages and 32 bulk soils.

### Bacterial DNA extraction, amplification and sequencing

DNA was extracted from all samples using DNeasy® PowerSoil® Pro Kits (QIAGEN®, Hilden, Germany) following the manufacturer’s protocol and stored at -20 °C. Concentrations and purity of the extractions were assessed using a Nanodrop2000™ spectrophotometer (Thermo Fischer Scientific™, Waltham, USA). DNA extraction yielded insufficient concentration and purity in 11 samples, which were therefore excluded from amplification. The amplification of full-length 16S rRNA gene (V1-V9 regions) was performed according to the PacBio® procedure and general best practices recommendations. Single-molecule Real-time long reads sequencing was performed at Gentyane Sequencing Platform (Clermont-Ferrand, France) with a PacBio Sequel II Sequencer (Pacific Biosciences®, Menlo Park, CA, USA). Amplicons were normalized at 5 ng/µL, pooled and purified with 1.3X SMRTbell cleanup beads. The SMRTbell library was prepared using a SMRTbell prep kit 3.0, following the “Preparing multiplexed amplicon libraries using SMRTbell prep kit 3.0” protocol. A Femto Pulse (Agilent Technologies®, Santa Clara, CA, USA) assay was used to assess the fragments size distribution. Pools were carried into the enzymatic reactions to remove the single-strand overhangs and to repair any damage that may be present on the DNA backbone. An A-tailing reaction followed by the overhang adapter ligation was conducted to generate the SMRTbell template. After nuclease treatment, pools were cleanup with 1.3X SMRTbell cleanup beads. The SMRTbell libraries were quality inspected and quantified on a Femto Pulse (Agilent Technologies®) and a Qubit fluorimeter with Qubit dsDNA HS reagent Assay kit (Life Technologies®). A ready-to-sequence SMRTbell Polymerase Complex was created using a Binding Kit 3.1 (PacBio®). The PacBio Sequel II instrument was programmed to load a 150pM library and sequenced in CCS mode on a PacBio SMRTcell 8M, with the Sequencing Plate 2.0 (Pacific Biosciences®, Menlo Park, CA, USA) with 0.5 hour of pre-extension time and acquiring one movie of 10 hours.

Four SMRTcells were used and the CCS reads were generated from raw data (i.e., subreads) with “ccs” and each sample was demultiplexed from each pool using “lima”. These were performed using “SMRT-Link v13.0” software (Pacific Biosciences®, Menlo Park, CA, USA). Two hundred and forty-nine samples were analyzed in a fastq file format. The DADA2 package (Callahan *et al.*, 2016) in R was used to perform taxonomic affiliations of the samples (see supplementary material). Primers were removed and strands orientation was conducted. Then, reads were filtered according to their sizes (between 1,000 and 1,600bp), followed by a dereplication and denoising steps to obtain the amplicon sequence variants (ASV, mean length = 1,449 bp). Taxonomic affiliations were done using the “silva_nr99_v138” database (Yilmaz *et al.*, 2014), and ASVs present in less than 2 samples were removed. The total number of ASVs retrieved was 19,747 (ccs reads data available in Genbank; SAMN45805597-SAMN45805845). Chloroplast, mitochondrial or eukaryote 16S rRNA sequences were removed, which resulted in 18,335 ASVs for 249 samples (see supplementary material, https://doi.org/10.5281/zenodo.18400087).

### Community ecology and statistical analyses

The statistical analyses were performed using RStudio software (version 4.3.0). To ensure data quality and that sampling of the different microhabitats was properly conducted, outlier samples from the initial set of 249 were identified using two approaches. Firstly, richness of the samples was calculated, and 62 outliers with aberrantly low or high richness were removed (likely due to technical issues that do not reflect true biological variation). Secondly, a Bray-Curtis dissimilarity matrix was constructed, and Nonmetric Multidimensional Scaling (NMDS) analysis was performed for each microbial community compartment using the “vegan” package (Oksanen *et al.*, 2022). The NMDS coordinates were used to calculate Mahalanobis distances between samples within each microhabitat. Outliers were defined as points with Mahalanobis distances exceeding the 99% Chi-squared threshold (Appendix 2), resulting in the removal of seven additional outliers and a final dataset of 180 samples. Finally, we removed ASVs representing less than 0.01% of total sequences to reduce computational load for phylogenetic inference, leading to a set of 1,590 ASVs. The same filtered dataset was then used for ecological and ecophylogenetic analyses, with minimal impact on results and biological interpretation (Appendix 3). The completeness of community coverage was confirmed using rarefaction curves (Schöler *et al.*, 2017). Shannon index (H) and species richness (S) were calculated using the *diversity* function in the “vegan” package. Analyses of the alpha diversity indexes were conducted using multiple Kruskal-Wallis test from the package “agricolae” (de Mendiburu, 2023). Community structure was analyzed and visualized using distance-based redundancy analysis with Bray-Curtis distances (db-RDA, capscale, “vegan”, 10,000 permutations; Oksanen *et al.*, 2022). This technique has proven to be useful when measuring multispecies responses to structured multifactorial designs (Legendre & Anderson, 1999). We assessed the homogeneity of multivariate dispersions by applying permutation tests to the residuals of the db-RDA model (Appendix 4). Differential abundance analyses were performed using “edgeR” (Robinson *et al.*, 2010). ASV count data were filtered to retain taxa with counts per million normalization > 1 in at least two samples and normalized by scaling each sample to the median library size. Differential abundance was assessed by contrasting bulk soil with each plant-associated microhabitat, and S1 to S3 within each plant-associated microhabitats (|log2FC| > 1, *p* < 0.05). Data and scripts required to reproduce the analyses are available at https://doi.org/10.5281/zenodo.18400087.

### Bacterial phylogenetic inference

In order to infer a bacterial phylogeny including ASVs we retrieved from our samples, we aligned 1,590 16S rRNA sequences using MUSCLE 3.6 (Edgar, 2004) and used Gblocks 0.91 (Castresana, 2000) to eliminate poorly aligned positions. The resulting matrix included 1,590 taxa and 1,195 characters (16S rRNA sequence matrix is available with the online version of the manuscript). Phylogenetic inferences were performed using BEAST 2.6.4 (Bouckaert *et al.*, 2019). The substitution model used was GTR +I+G (Yang, 1994); three runs of 100 million chains were performed, sampling every 5,000 generations. We checked for convergence and viewed statistics for each run using TRACER 1.7.2 (Rambaut *et al.*, 2018). After combining the three runs and removing 40% of burn-in samples, we summarized the results using a maximum clade credibility (MCC) tree.

We computed the Rao’s quadratic entropy (FD_Q,_ Rao, 1982, “SYNCA”, Debastiani & Pillar, 2012), a weighted measure of diversity integrating both phylogenetic distances and relative abundances of bacterial taxa, to assess the effects of host-filtering on phylogenetic diversity. Temporal phylogenetic dispersion model of bacterial communities was estimated by calculating the empirical dispersion parameter (*D*) for each plant-associated microhabitats, following the method developed by Darcy *et al.* (2020). The dispersion parameter tracks how newly recruited taxa relate phylogenetically to previously observed taxa over time, allowing to test whether new recruits are more closely or more distantly related to earlier community members than expected by chance (Darcy *et al.*, 2020). All above analyses were performed using the MCC tree.

## Results

### Bacterial community composition and structure

Bacterial communities were characterized from a total of 180 samples. To ensure a good coverage of the bacterial communities, we generated rarefaction curves using the full set of 18,335 ASVs, illustrating the increase in observed ASVs as a function of sequencing depth (Appendix 5). All samples reached an asymptote, indicating that the bacterial communities were fully sampled in all microhabitats and that rarefaction was unnecessary. Community diversity did not differ significantly among the four *Pisum* accessions, and although community structure varied significantly, the explained variance was low (< 5%, Appendix 6). Therefore, all genotypes were combined for subsequent analyses. Bacterial community diversity varied across the different plant-associated microhabitats (Figure 1A, B). Specific richness was the highest in the bulk soil (bulk, median S = 547), followed by the rhizosphere (R, S = 399) and rhizoplane (Rp, S = 389), with the lowest richness observed in the endosphere (E, S = 70; Figure 1A, *p* < 0.001). The Shannon diversity index significantly decreased across the microhabitats, from 5.73 in bulk to 5.42 in R, 5.14 in Rp, and 3.62 in E (Figure 1B, *p* < 0.001). Although specific richness and Shannon diversity were not significantly influenced by growth stages, it is noteworthy that the initial community exhibited the highest diversity, with S = 660 and H = 6.05 in the initial bulk soil (i.e., soil sampled before sowing) (Appendix 7).

**Figure 1.**
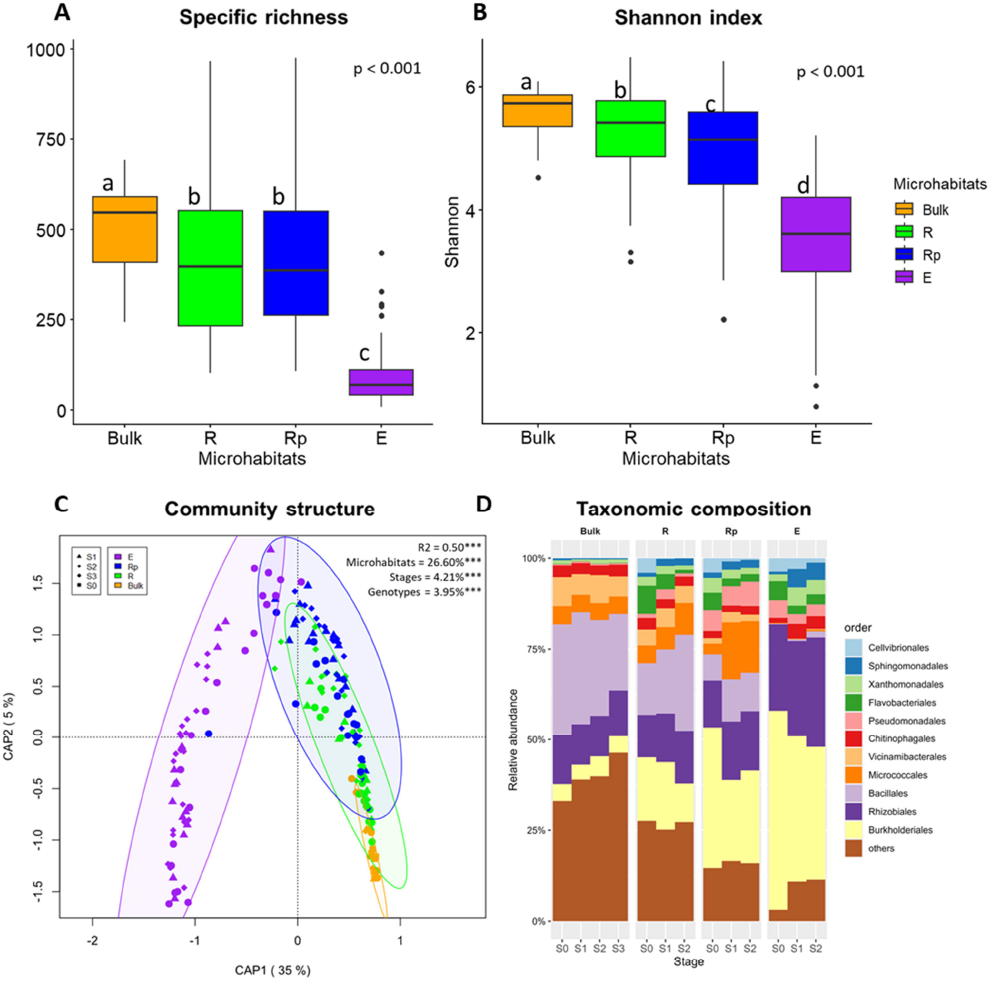
Bacterial community diversity, structure and composition. **Panel A-8:** Specific richness (A) and Shannon diversity index (BJ of bacterial communities across bulk soil and the different plant-associated microhabitats. Multiple Kruskal-Wallis comparisons. **Panel C:** Broy-Curtis distance-based redundancy analysis (similar to Constrained Analysis of Principal Coordinates). The model was tested using PERMANOVA with 10,000 permutations. The R-squared value indicates the percentage of variance explained by the model, including the three variables (microhabitat, growth stages and host genotype) and their interactions. Note that for figure clarity, host genotypes are not displayed. **Panel D:** Average taxonomic composition of bacterial communities based on relative abundance at the Order level across bulk soil, plant-associated microhabitats and growth stages. Only the eleven most abundant Orders are shown, with the remaining Orders grouped together and labeled as “others”. Note that data from all host genotypes were compiled. (N_totof_ = 180, N_Bu_/k = 21, N_R_ = 46, N_Rp_ = 56, N_E_= 57)

Bacterial communities’ structure differs across plant-associated microhabitats, growth stages and host *Pisum* genotypes (R^2^ = 0.50, *p* < 0.001, Figure 1C). Changes in structure were best explained by microhabitats (26.60%, *p* < 0.001, Figure 1C). There was no overlap of 95% CI ellipses between microhabitats that were not in close physical proximity (Figure 1C). The highest overlap in 95% CI was observed between R and Rp, while the overlaps with the other microhabitats were small. Growth stages had a lower but significant effect on bacterial communities’ structure (4.21%, *p* < 0.001; Figure 1C). The host genotype had a significant but marginal influence on the associated bacterial communities (3.95%, *p* < 0.001; Figure 1C). Time had no significant effect on the bacterial communities in the bulk soil (*p* = 0.15; Appendix 8A). In contrast, growth stages had a significant and increasingly pronounced effect on the bacterial communities in the plant-associated microhabitats, explaining respectively 9.31%, 10.92%, and 13.27% of the variance in R, Rp and E (*p* < 0.001; Appendix 8B, C, D).

The taxonomic composition of bacterial communities varied across plant-associated microhabitats (Figure 1D, Appendix 9A, B, C). Bulk soil communities were the most heterogeneous, with low-abundance orders (“others”) accounting for 39.6% of relative abundance, alongside Bacillales (27.2%), Rhizobiales (12.2%), Vicinamibacterales (6.7%), Micrococcales (4.8%) and Burkholderiales (4.7%) (Figure 1D). In the rhizosphere, community composition remained broadly similar to bulk soil but showed an enrichment in Burkholderiales (15.6%), mainly driven by ASVs affiliated to *Massilia, Rhizobacter, Ramlibacter* and *Noviherbaspirillum* (Appendix 10A), together with a decrease in Bacillales (19.5%). In the rhizoplane, communities shifted toward a dominance of Burkholderiales (28.9%), Micrococcales (11%) and Pseudomonadales (5.9%), driven by *Delftia, Massilia, Paeniglutamicibacter* and several *Pseudomonas* species (Appendix 10B), while Bacillales (9.8%) and Vicinamibacterales (2%) decreased. In the endosphere, communities were dominated by Burkholderiales (43.8%) and Rhizobiales (26.8%), with Bacillales, Micrococcales and Vicinamibacterales nearly absent. The highest increase in the endosphere is for *Rhizobium leguminosarum* (Appendix 10C). Notably, several orders that were nearly absent in bulk soil (<1%), including Pseudomonadales, Flavobacteriales, Xanthomonadales, Sphingomonadales and Cellvibrionales, progressively increased in relative abundance across plant-associated microhabitats (Figure 1D).

Community composition within plant-associated microhabitats also varied across growth stages (S1 to S3), whereas no significant differences were observed in bulk soil (Figure 1D, Appendix 9D, E, F, G). In the rhizosphere, Bacillales (14.3 to 26.5%), Rhizobiales (11.7 to 14.7%) and Micrococcales (4.9 to 8.7%) increased, driven mainly by *Bacillus* species, *Paeniglutamicibacter sulfureus* and *Paenarthrobacter aurescens* (Appendix 10D), while Burkholderiales (17.6 to 10.6%), Flavobacteriales (7.8 to 0.9%) and Cellvibrionales (4.0 to 0.1%) declined. In the rhizoplane, Micrococcales (2.9 to 14.2%) and Pseudomonadales (5.8 to 6.7%) increased, driven primarily by *Paeniglutamicibacter sulfureus* and *Pseudomonas* species (Appendix 10E), whereas Burkholderiales (38.8 to 25.5%), Cellvibrionales (3.9 to 0.4%), Flavobacteriales (4.7 to 2.1%) and Xanthomonadales (4.1 to 1.7%) decreased. In the endosphere, the only marked increase was observed for Rhizobiales (22.8 to 31.8%), mainly driven by *Rhizobium leguminosarum* (Appendix 10F), while Burkholderiales (54.6 to 35.2%), Cellvibrionales (3.7 to 1.2%) and Pseudomonadales (4.7 to 2.9%) declined.

### Temporal and spatial recruitment in a phylogenetic context

Using our phylogenetic tree of 1,590 bacterial ASVs, we examined the phylogenetic diversity of bacterial communities across the different microhabitats and calculated the phylogenetic dispersion parameter (*D*) to assess the phylogenetic patterns of bacterial recruitment throughout growth stages in each plant-associated microhabitat. Negative *D* values indicate underdispersion, corresponding to the preferential recruitment of phylogenetically close taxa from the established community over time, whereas positive values indicate overdispersion, reflecting the recruitment of phylogenetically more distant taxa. Phylogenetic diversity was the highest in the bulk with FD_Q_ = 0.67 (*p* < 0.001, Figure 2A) and progressively decreased across the plant-associated compartments with FD_Q_ = 0.65 in R, FD_Q_ = 0.60 in Rp and FD_Q_ = 0.44 in E (*p* < 0.001, Figure 2A). The phylogenetic diversity significantly varied across growth stages only within the endosphere, rising from FD_Q_ = 0.40 at S1 to FD_Q_ = 0.44 at S2 and FD_Q_ = 0.46 at S3 (*p* < 0.01, data not shown). The phylogenetic dispersion parameter differs across plant-associated microhabitats (Figure 2B). Bacterial communities in R exhibited overdispersion pattern with *D* > 0 (*D* = 0.34, *p* < 0.05, Figure 2B). In Rp, *D* was lower than in R (*p* < 0.1) and not different to *D* = 0, indicating a neutral phylogenetic dispersion pattern (*D* = 0.03, *p* > 0.05, Figure 2B). Communities in E showed underdispersion, with *D* = -0.40, significantly lower than in Rp and different from the neutral model (*p* < 0.05, Figure 2B), indicating that closely related ASVs have more probabilities of being recruited throughout growth stages.

**Figure 2.**
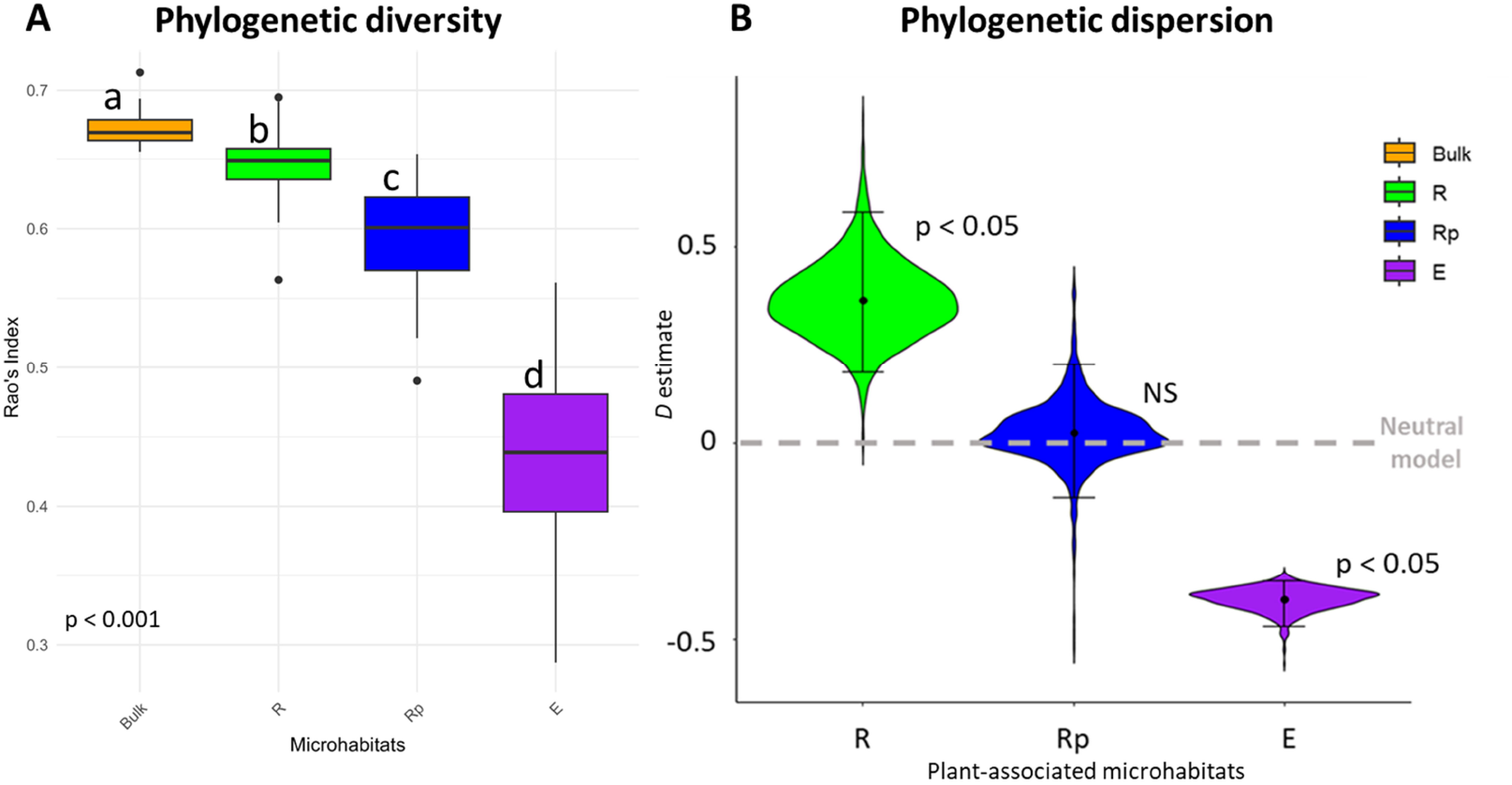
Phylogenetic diversity and dispersion parameter of bacterial communities across the different microhabitats. **Panel A:** Abundance weighted phylogenetic diversity (Rao’s quadratic entropy index, FD_0_) (Rao, 1982) of bacterial communities across the different microhabitats. Multiple Kruskal-Wal/is test, p < 0.001, N = 180. **Panel B:** Phylogenetic dispersion parameter (D) (Darcy et al., 2020) of bacterial communities across the different plant-associated microhabitats. D > 0 indicates overdispersion, D = 0 is the neutral model and D < 0 indicates underdispersion. N = 159. The p-values indicate significant differences in D values to neutral model, rather than differences between microhabitats.

## Discussion

### Microbial community assembly

Bacterial communities were specific to the different microhabitats. These offer distinct ecological niches, which is reflected by the significant changes in abundance, composition, and structure of bacterial communities. The highest diversity was observed in the bulk soil and gradually decreased from the rhizosphere to the endosphere. Plant-associated microhabitats act as ecological filters along the spatial gradient, where communities from the bulk soil are shaped in the rhizosphere, which are further influenced by the rhizoplane, ultimately resulting in only a small part of these communities being present in the endosphere. Community composition exhibited a clear successional shift from bulk soil to the endosphere, marked by a progressive enrichment of several orders nearly absent in bulk soil, including Pseudomonadales, Flavobacteriales, Xanthomonadales, Sphingomonadales, and Cellvibrionales. Notably, Burkholderiales became dominant along the gradient, while orders such as Bacillales and Vicinamibacterales decreased. Within the Burkholderiales, potential PGP genera such as *Massilia* (Dai *et al.*, 2020; Angot *et al.*, 2025) and *Noviherbaspirillum* (Barros *et al.*, 2022) drove these shifts. Within Pseudomonadales, *Pseudomonas* species, which are common in the endosphere and nodules of legumes (Mayhood & Mirza, 2021; Hossain *et al.*, 2023) and may influence the outcome of root-nodule symbiosis (Crosbie *et al.*, 2022) drove the signal. The increase in Rhizobiales in the endosphere was mainly driven by *Rhizobium leguminosarum*, likely due to the spatial proximity with root nodules.

Host growth stages also influenced the structure and composition of bacterial communities within plant-associated microhabitats, while bulk soil communities remained stable. In the rhizosphere, we observed an increase in Bacillales driven by potential PGP *Bacillus species* (El-Esawi *et al.*, 2018; Blake *et al.*, 2021), and Micrococcales, mainly driven by *Paeniglutamicibacter sulfureus*, alongside a decline in Burkholderiales, Flavobacteriales, and Cellvibrionales. The rhizoplane exhibited a pronounced increase in Micrococcales, primarily due to *Paeniglutamicibacter sulfureus*, and a slight increase in Pseudomonadales (*Pseudomonas spp.*), while Burkholderiales and several other orders decreased. This marked enrichment of *Paeniglutamicibacter* at later stages is notable. Although its direct role in plant interactions is unknown, its capacity for hydrocarbon degradation in soils (Brzeszcz *et al.*, 2024) suggests a potential response to root metabolites, given the structural similarities between hydrocarbons and many plant secondary metabolites (Singer, 2006; Rohrbacher & St-Arnaud, 2016). In the endosphere, the most significant temporal change was a further increase in Rhizobiales, dominated by *Rhizobium leguminosarum*, consistent with the hypothesis of dispersion from root nodules, whose abundance rises with plant development (Mortier *et al.*, 2012).

The stability of bulk soil communities, compared to plant-associated microhabitats, highlights the importance of host’s influence on the temporal dynamics of bacterial communities. Even if further work is necessary to understand the mechanistic processes driving plant-bacterial recruitment throughout plant development, numerous studies have illustrated the influence of plant growth stages on the non-symbiotic microbial communities (Qiao *et al.*, 2017; Dai *et al.*, 2019; Larrouy *et al.*, 2023). These variations seem to be tied to shifts in the plant’s requirements throughout its development, as well as to changes in the quality and quantity of rhizodeposits produced by the host (Hütsch et al., 2002; Houlden et al., 2008; Chaparro et al., 2013). Indeed, rhizodeposition is dependent on plant carbon allocation (Shamoot et al., 1968) which varies through growth stages (Morin et al., 2022). Carbon allocation in the roots is important during the vegetative stage (Voisin et al., 2003b). At the flowering stage, the concentration of carbon increases to supply shoot apical meristem and induces floral transition (Cho et al., 2018). During the reproductive stage, carbon is remobilized from the leaves, stem and roots to fill the seeds (Weber et al., 1997). Because the microorganisms associated with *Pisum* spp. can enhance its nitrogen acquisition, the dynamics of the rhizosphere microbiota are linked to the plant’s nitrogen needs, which vary at different growth stages (Du et al., 2023). During the vegetative phase, nitrogen requirements are significant to ensure plant growth and to build reserves in the above-ground parts (Voisin et al., 2007). During seed filling stages, nitrogen acquisition is reduced and the plant uses its nitrogen stocks to fill the seeds (Voisin et al., 2003a). Both carbon and nitrogen fluxes variations impact the rhizosphere microbiota; however, the influence of developmental stages on the recruitment of microbial communities remains uncertain (Biget et al., 2024), especially given the potential confounding effect of the time since sowing (Dibner et al., 2021).

Our findings reveal that plants host diverse bacterial communities that vary across the microhabitats they provide and throughout their growth stages. These shifts in community composition and structure reflect the plant’s capacity to act as an ecological filter, creating distinct niches that select for particular bacterial taxa. At the same time, microorganisms also contribute to shaping their environments, feeding back into the construction and modification of plant-associated microhabitats. Given the complexity of integrating spatial and temporal factors in host-microbiota systems, moving toward an eco-evolutionary framework appears particularly valuable. Such an approach would help disentangle the mechanisms driving community recruitment, persistence, and changes across plant development.

### Phylogenetic constraints on bacterial community assembly

Our bacterial phylogeny, based on full-length 16S rRNA gene sequencing of 1,590 ASVs, is largely congruent with recent genomic references (Zhu *et al.*, (2019), Appendix 11). All Bacterial clades were retrieved (Appendix 11). Proteobacteria (with Gamma- and Alphaproteobacteria as sister groups), Desulfobacterota (sister to Proteobacteria), and Bacteroidota. Firmicutes, Actinobacteria, Chloroflexi, and Cyanobacteria are also monophyletic. The placement of Acidobacteriota differs, possibly due to our denser sampling (128/1,590 vs 56/10,575 taxa in Zhu et al. (2019)). Planctomycetota, not sampled in Zhu et al. (2019), are placed as sister to Thermotogae, which is sister to Cyanobacteria in Zhu et al. (2019), supporting our topology (Rensink *et al.*, 2020). These results suggest that, although recombination and horizontal gene transfer processes affecting the 16S rRNA gene can weaken the concordance of 16S rRNA phylogenies (Hassler *et al.*, 2022), using full-length sequencing enables the construction of phylogenies that are congruent with phylogenomic inferences. In addition, 16S rRNA gene phylogenies were shown to reflect the ecological and functional traits of soil bacterial communities (Borenstein *et al.*, 2008). The phylogenetic diversity (PD) varied across microhabitats, with the highest PD being observed in the bulk soil, followed by a gradual decrease across the rhizosphere, rhizoplane, and endosphere. Host filtering does not randomly reduce bacterial diversity but rather selects for specific, phylogenetically related lineages, likely by intensifying niche constraints. Goldford *et al.* (2018), using ex situ cultivation of microbial communities from different habitats, showed that communities were assembled at family-level, suggesting that this pattern may reflect functional selection linked to competitive advantages. Our results support the prediction that phylogenetically conserved set of traits allow specific bacteria lineages to pass through ecological constraints imposed by the host plants (Aguirre de Cárcer, 2019). This suggests that selective recruitment is correlated with bacterial functions required for plant development, as previously reported by Mougel *et al.* (2006). In addition, the temporal phylogenetic pattern of community assembly varied across the plant-associated microhabitats. The model we used considers the likelihood that newly recruited taxa are similar to closely related taxa previously recruited (Darcy *et al.*, 2020). Overdispersion observed in the rhizosphere suggests that interspecific competition among closely related bacteria plays a crucial role in shaping the community (Nemergut *et al.*, 2013). Closely related taxa, which share similar niches, are competitively excluded, leading to phylogenetic overdispersion of the bacterial community. The rhizosphere, as an open and nutrient-rich microhabitat, could provide a diverse array of ecological niches. These niches can support the coexistence of distantly related bacteria, probably with distinct ecological functions. Conversely, the underdispersion pattern observed in the endosphere indicates the opposite process: the bacterial community is not mainly structured by competition among closely related bacteria but rather by strong host-filtering (Mayfield & Levine, 2010), as previously observed for mycorrhizal fungal community assembly (Frew *et al.*, 2023). The endosphere, as a highly selective and closed microhabitat, imposes significant host-mediated environmental constraints that can be physical (barriers of root cells) or chemical (plant-derived chemical compounds; Lumibao *et al.* (2020)). It shows that closely related bacteria are able to coexist in a community (Webb *et al.*, 2002), with specific functional traits, often conserved among phylogenetically related taxa. The pattern of phylogenetic underdispersion in the endosphere is consistent with ecological selection acting on phylogenetically conserved traits (Webb *et al.*, 2002; Gerhold *et al.*, 2015). If the traits being selected for are horizontally transferred between bacteria lineages with no connection with their relatedness, we would not observe any phylogenetic signature (Darcy *et al.*, 2020). Several studies have demonstrated that plant hosts actively select for specific bacterial communities or strains. (Bertin *et al.*, 2003; Bulgarelli *et al.*, 2013; Sasse *et al.*, 2018; Bourion *et al.*, 2018; Xiong *et al.*, 2021). Over evolutionary timescales, this persistent host-mediated ecological filtering could have driven the evolutionary history of plant-associated bacterial communities. Such reciprocal interactions, which impact the fitness of both partners, could be a keystone in the evolutionary trajectory of land plants and a mark of coevolution (Fitzpatrick et al., 2018).

Altogether, our results illustrate the strong influence of plants on their associated soil bacterial communities, shaping both their ecological and evolutionary dynamics across spatial and temporal scales. Bacterial communities were highly specific to underground plant-associated microhabitats, and their assembly was phylogenetically structured, with the host filtering taxa according to their evolutionary relatedness. However, investigating the heritability of bacterial communities is crucial for understanding whether these associations are shaped by (i) horizontal transfer from the environment to plant-associated microhabitats, (ii) plant-specific traits, or (iii) vertical transmission across plant generations. A limitation of this study is the absence of direct measurements of plant physiology and root exudates. Consequently, while we document clear spatio-temporal patterns in community assembly, it cannot be directly linked to bacterial plant growth promotion or plant recruitment mechanisms. Future work integrating host phenotyping, exudate profiling, and microbial functional potential is essential for a mechanistic understanding of the plant-bacteria interactions and their evolutionary outcomes.

## Supporting information

Appencides

## Acknowledgments

The authors thank the 4PMI greenhouse platform team at INRAe Dijon for help in designing the experiment and the ECP team (LEGae, INRAE Dijon) for valuable assistance in the laboratory. The authors wish to thank Nadine Praeg and Mathieu Barret for critically evaluating and providing very valuable comments that improved significantly the manuscript. This research was funded by INRAE and the Agence National de la Recherche (ANR, France).

