## Supplementary material for "Ecophylogenetic patterns of rhizosphere bacterial community assembly in *Pisum* spp. (Fabaceae, Fabeae) reveal strong plant-mediated ecological filtering": Appencides

Horizon : Soil
Depth (cm) : 0-30
N-NO3 (mg/kg) : 34.4
N-NH4 (mg/kg) : 0.20
Water (%) : 1.8%
Density : 1.35
N-NO3 (kg/ha) : 139.3
N-NH4 (kg/ha) : 0.8
N total (kg/ha) : 140.1

***Appendix 1:*** *Soil chemical analyses performed by SADEF Agronomy & Environment laboratory. Soil nitrate ($N–NO_3^-$) and ammonium ($N–NH_4^+$) were determined using KCl extraction according to ISO/TS 14256-2, followed by colorimetric quantification. Soil water content was assessed gravimetrically according to ISO 11465. Bulk density was determined according to ISO 11272.*

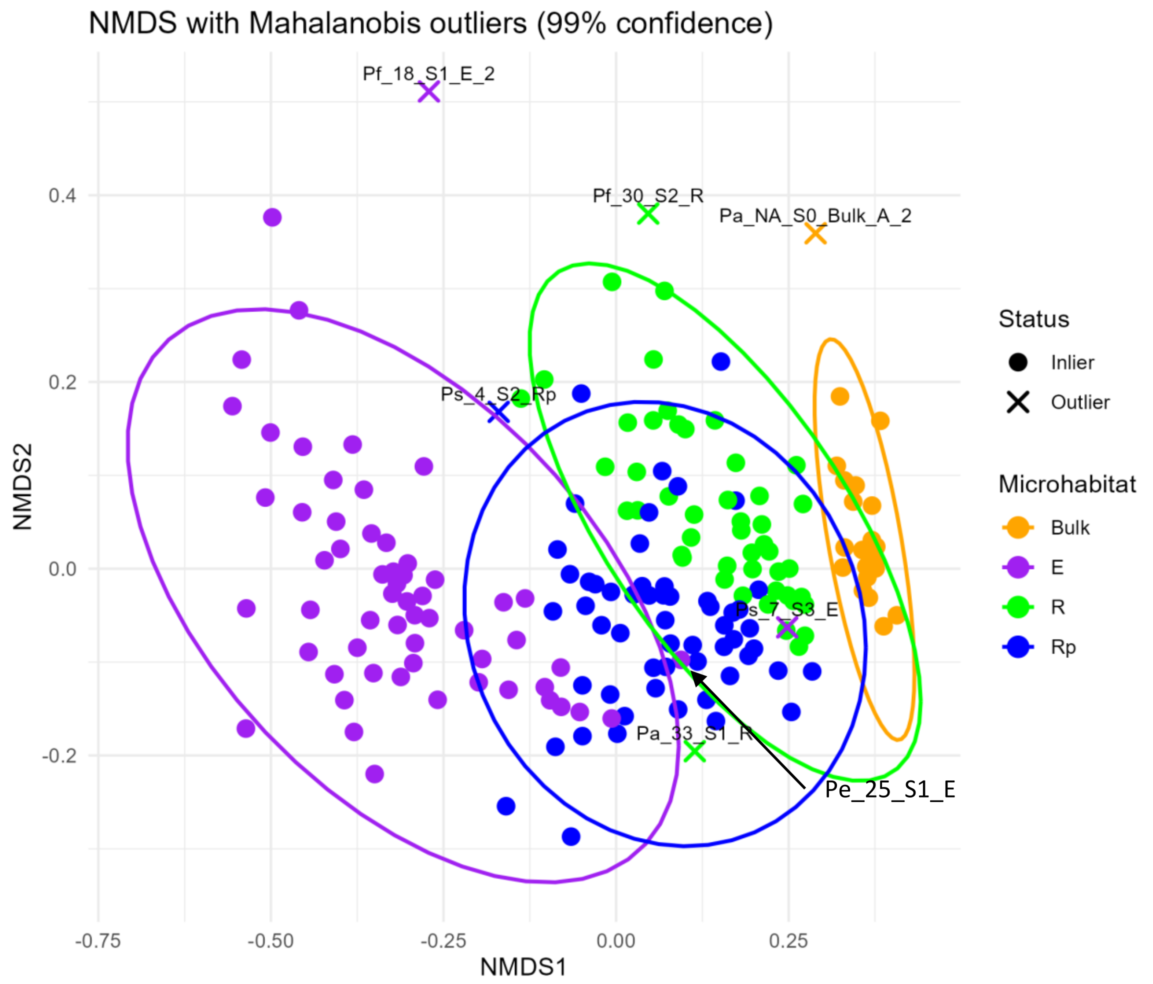


| Outliers | R | Rp | E | Bulk | Total |
|---|---|---|---|---|---|
| S0 | | | | 3 | 3 |
| S1 | 15 | 7 | 11 | 4 | 37 |
| S2 | 7 | 6 | 3 | 1 | 17 |
| S3 | 3 | 4 | 2 | 3 | 12 |
| Total | 25 | 17 | 16 | 11 | 69 |

***Appendix 2: Panel A:*** *Bray-Curtis distance based Nonmetric Multidimensional Scaling (NMDS) analysis and detection of outliers using Mahalanobis distances (Mahalanobis, 1930). The NMDS coordinates were used to calculate Mahalanobis distances within each microhabitat. Outliers defined as points with Mahalanobis distances exceeding 99% Chi-squared threshold. Sample "Pe_25_S1_E" was not detected as an outlier by this analysis but was manually removed due to aberrant richness and proximity to the threshold.*
***Panel B:*** *Distribution of outlier samples across the different microhabitats and growth stages. Outliers were detected based on the richness and structure of the bacterial communities.*

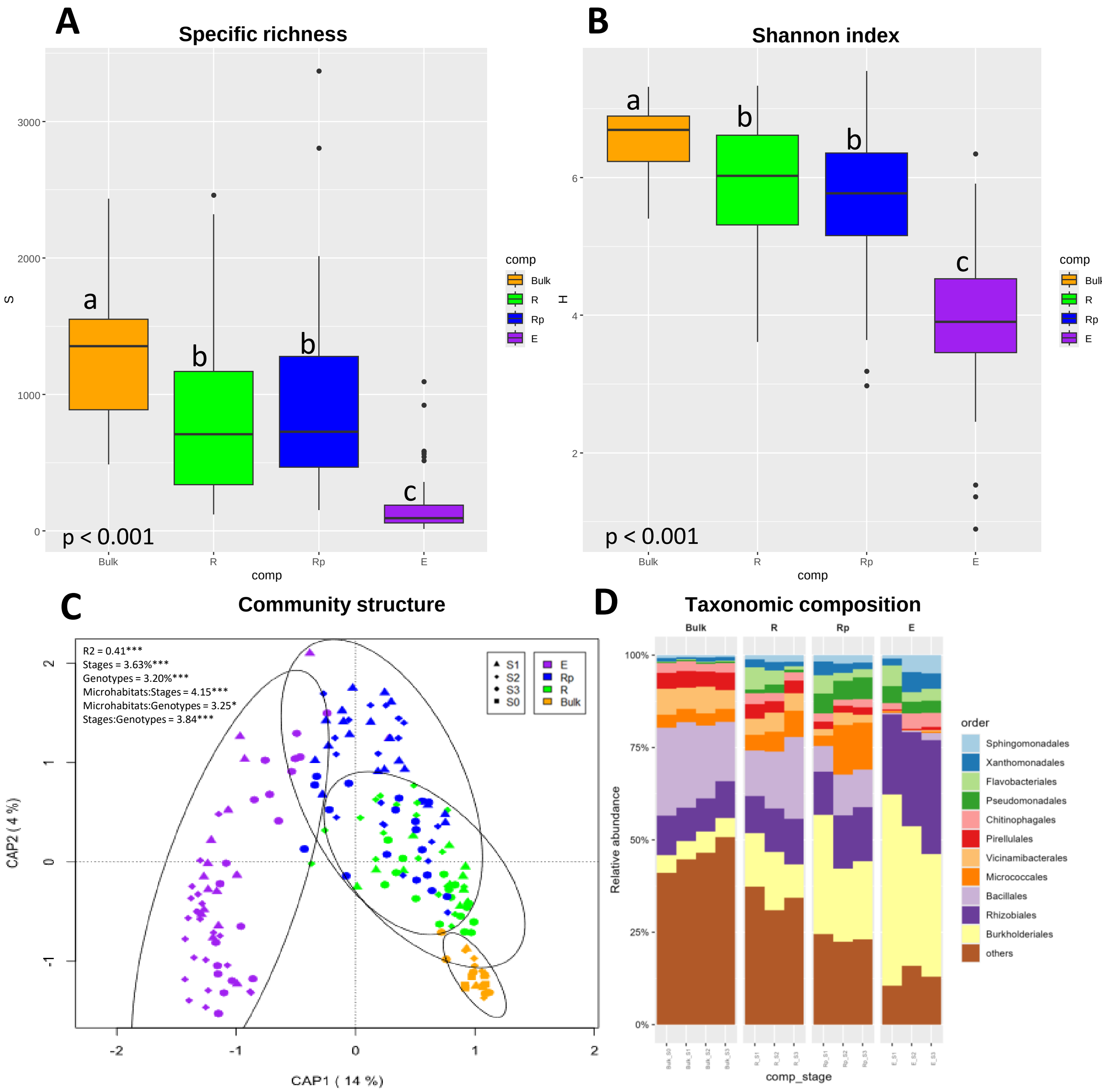


***Appendix 3:*** *Bacterial community diversity, structure and composition analyses using the total set of 18,335 ASVs, previous to filtering at 0.01% of relative abundance.* ***Panel A-B:*** *Specific richness (A) and Shannon diversity index (B) of bacterial communities across bulk soil and the different plant-associated microhabitats. Multiple Kruskal-Wallis comparisons.* ***Panel C:*** *Bray-Curtis distance-based redundancy analysis (similar to Constrained Analysis of Principal Coordinates). The model was tested using PERMANOVA with 10,000 permutations. The R-squared value indicates the percentage of variance explained by the model, including the three variables (microhabitat, growth stages and host genotype) and their interactions. Note that for figure clarity, host genotypes are not displayed.* ***Panel D:*** *Average taxonomic composition of bacterial communities based on relative abundance at the Order level across bulk soil, plant-associated microhabitats and growth stages. Only the eleven most abundant Orders are shown, with the remaining Orders grouped together and labeled as "others". Note that data from all host genotypes were compiled. ($N_{total}$ = 180, $N_{Bulk}$ = 21, $N_R$ = 46, $N_{Rp}$ = 56, $N_E$ = 57)*

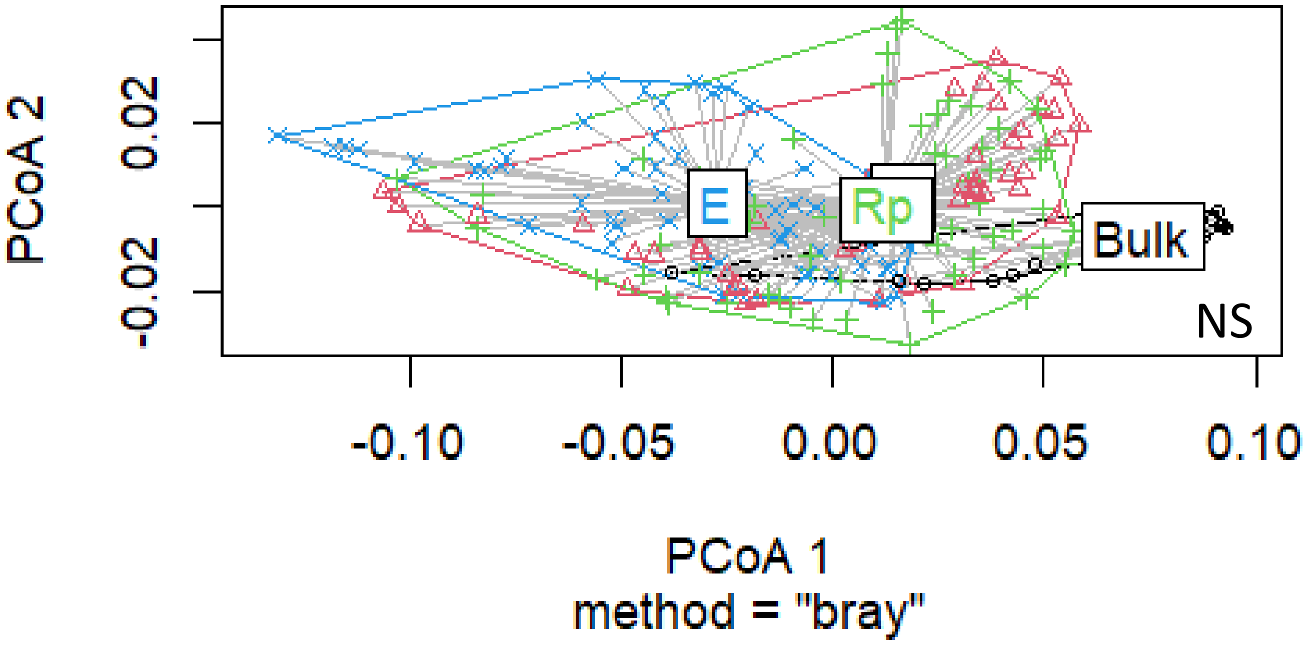


***Appendix 4:*** *Multivariate dispersion of residuals from the db-RDA model across microhabitats. Distances among residuals were calculated using Bray-Curtis dissimilarities, and group centroids were compared using betadisper. No significant differences in dispersion were detected among microhabitats (permutation test, p > 0.05), indicating homogeneous multivariate variance across groups.*

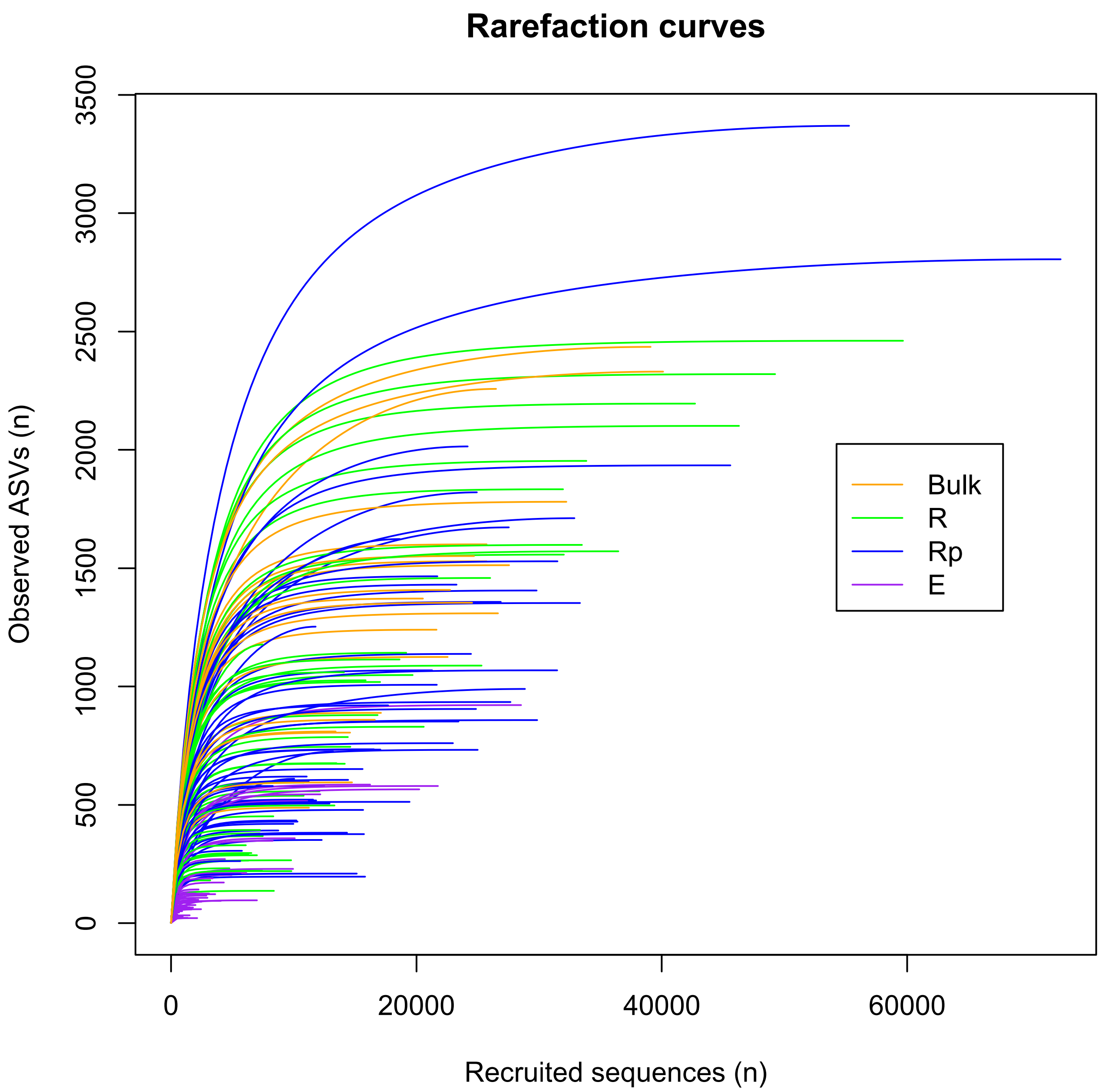


***Appendix 5:*** *Rarefaction curves generated prior to ASVs filtering (18.335 ASVs). Line colors indicates the microhabitat of the community.  N = 180.*

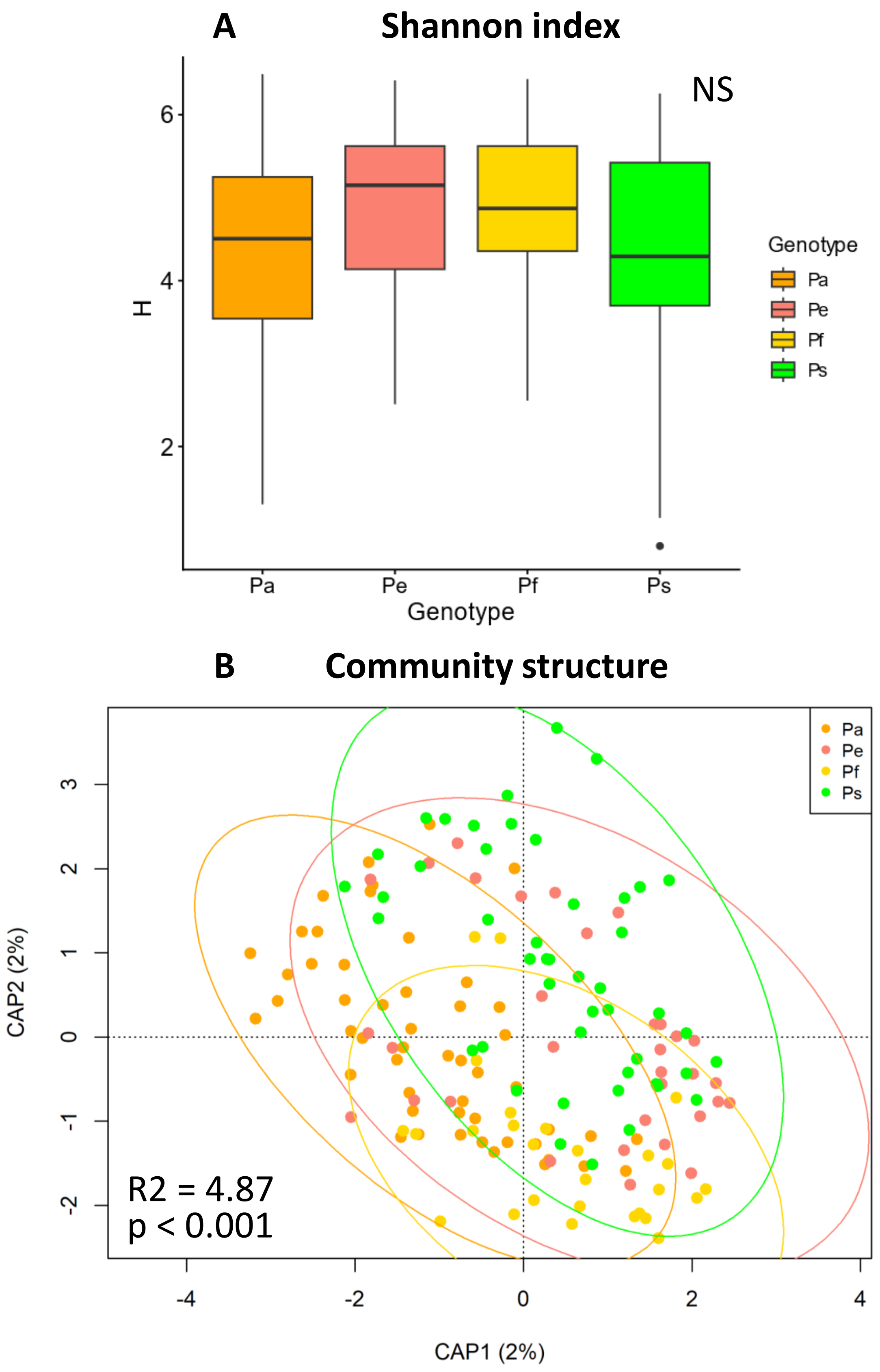


***Appendix 6:*** *Shannon index diversity and structure of bacterial communities across host genotypes.* ***Panel A:*** *Shannon index of bacterial communities across host genotypes (all microhabitats and stages considered). NS = not significant.* ***Panel B:*** *Structure of bacterial communities across host genotypes (all microhabitats and stages considered). R2 being the variance explained by the host genotypes.*

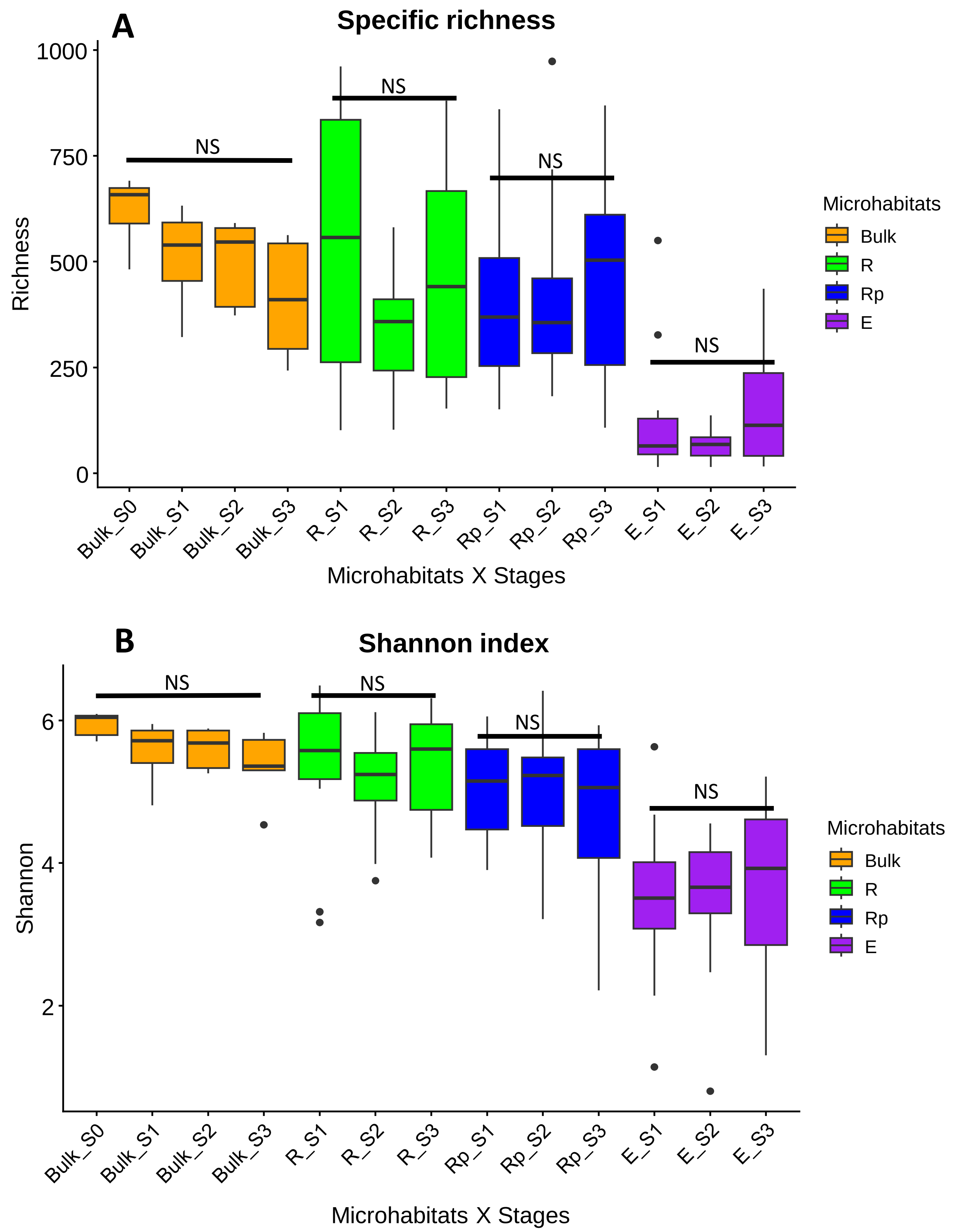


***Appendix 7:*** *Specific richness (A) and Shannon diversity index (B) of bacterial communities across bulk soil and the different plant-associated microhabitats through growth stages. Multiple Kruskal-Wallis comparisons within microhabitats. NS indicates non-significant differences. ($N_{total}$ = 180, $N_{Bulk}$ = 21, $N_R$ = 46, $N_{Rp}$ = 56, $N_E$ = 57)*

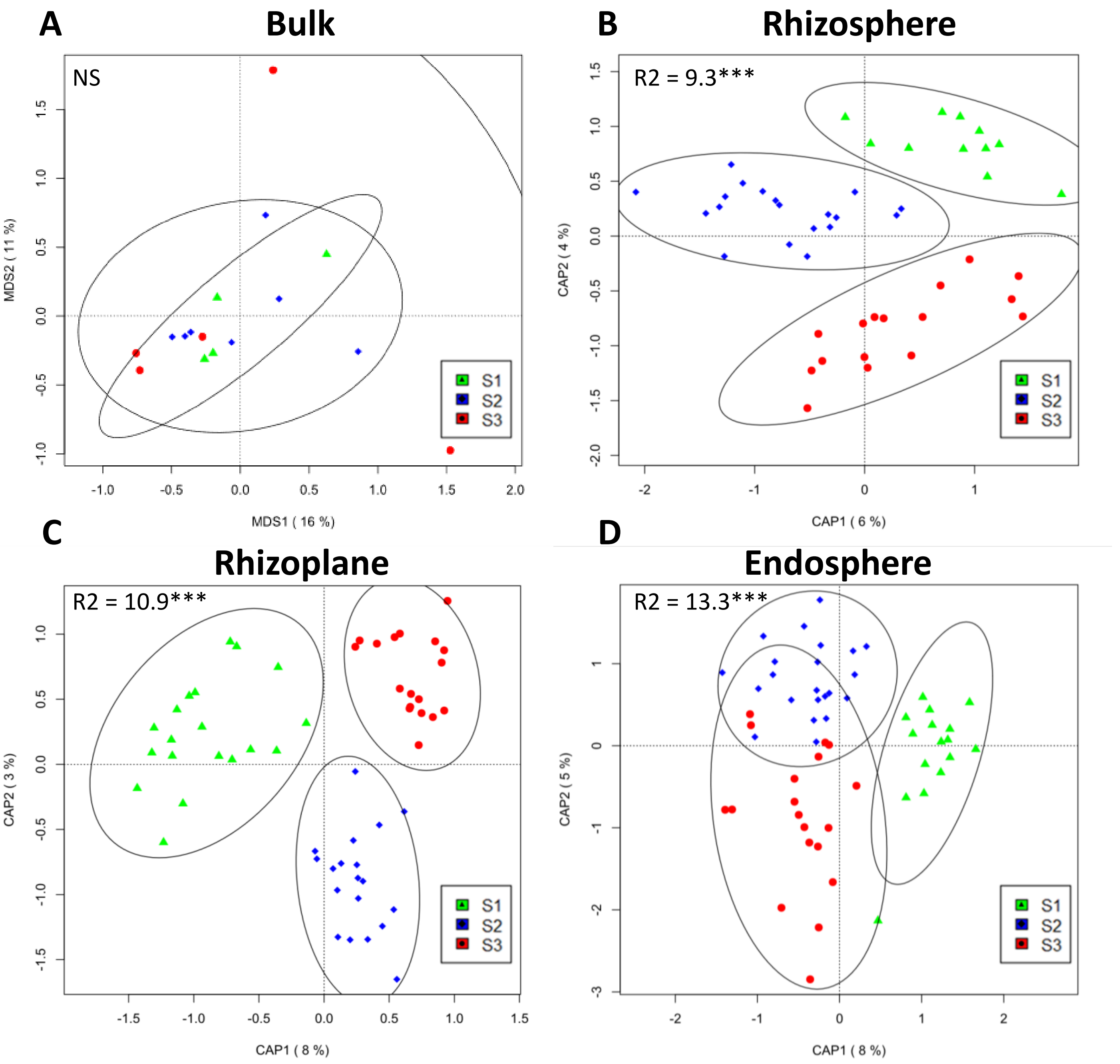


***Appendix 8:*** *Bray-Curtis distance-based redundancy analysis (similar to Constrained Analysis of Principal Coordinates, CAP) was performed on bacterial communities across growth stages for each microhabitat separately. The models were tested using PERMANOVA with 10,000 permutations. The R-squared value represents the percentage of variance explained by the model. When significant, constrained axes (CAP) are displayed; otherwise, multidimensional scaling (MDS) is shown. Significance code: NS p > 0.05; * p < 0.05; ** p < 0.01; *** p < 0.001.*

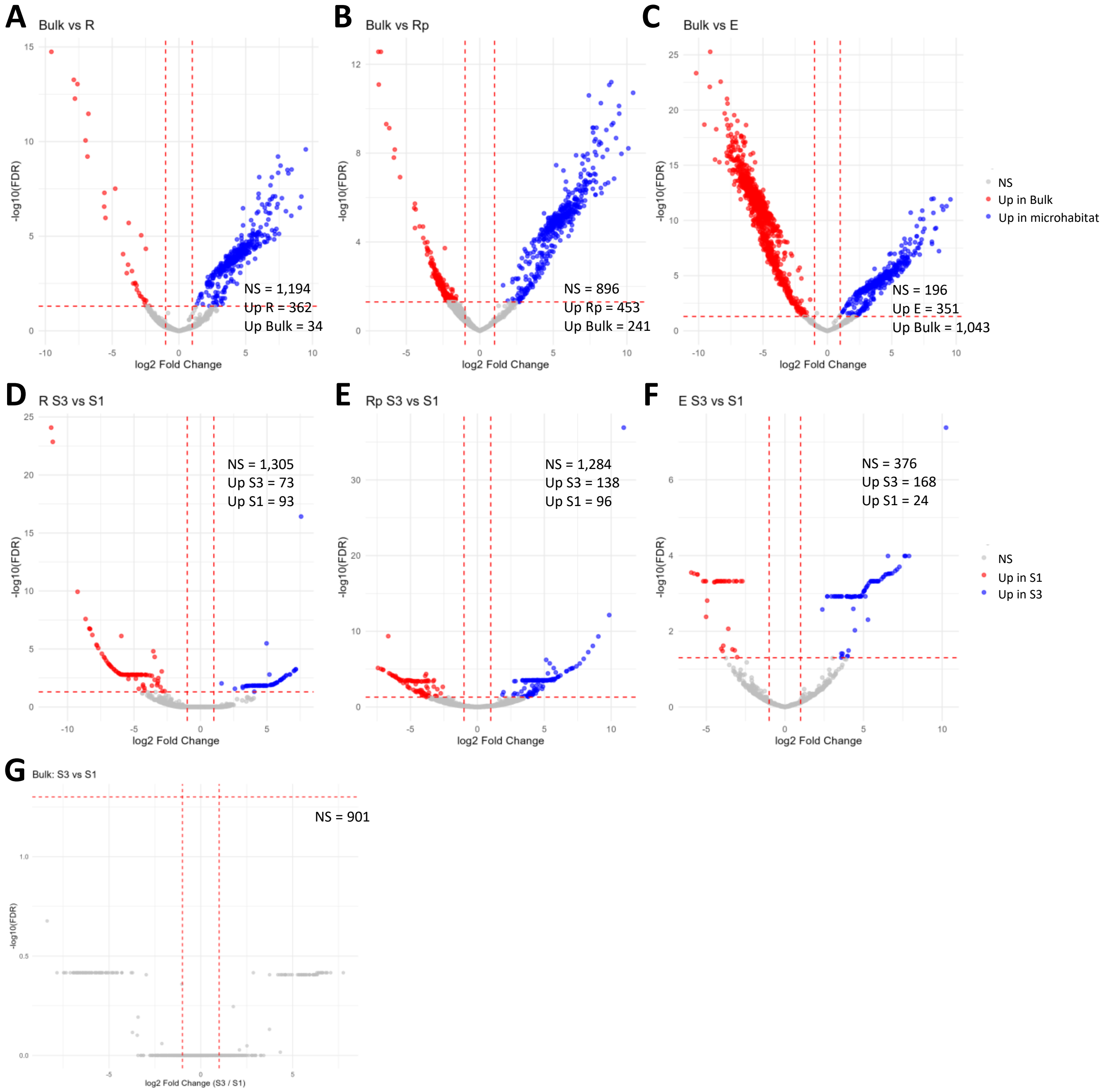


***Appendix 9:*** *Volcano plots showing differential abundance of ASVs between Bulk and plant-associated microhabitats (A, B, C), or between stage 1 and stage 3 within each microhabitats (D, E, F, G). The x-axis indicates the $log_2$ fold change between conditions, while the y-axis shows the $-log_{10}$ of the adjusted p-value.*

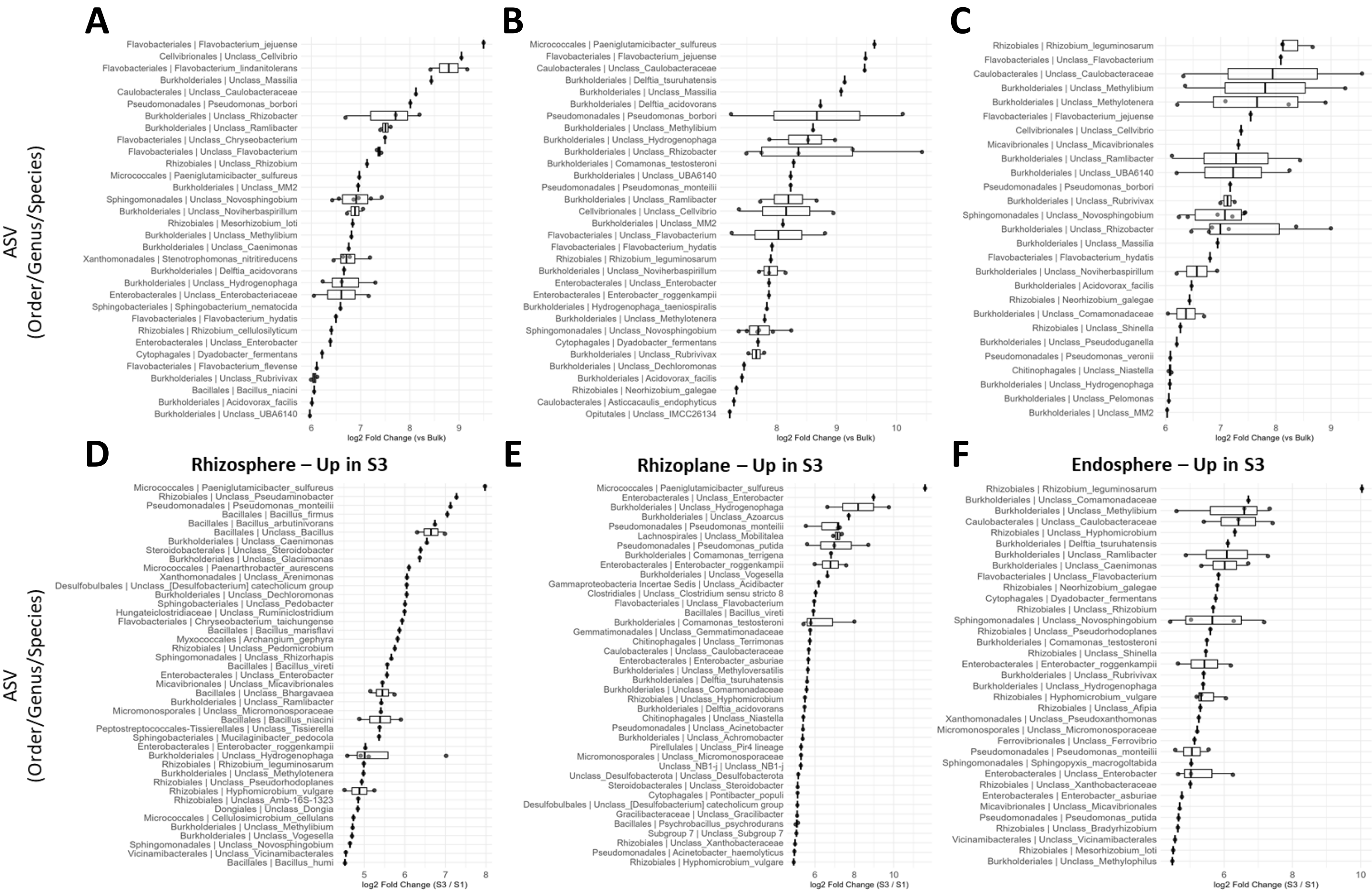


***Appendix 10:*** *Top 50 ASVs significantly enriched in microhabitats compared to Bulk soil (A, B, C) or at stage 3 compared to stage 1 within each plant-associated microhabitats. Each plot shows the 50 ASVs with the highest* $log_2$ *fold change (logFC, edgeR). Points represent individual ASVs, with Order, Genus and Species indicated on the y-axis.*

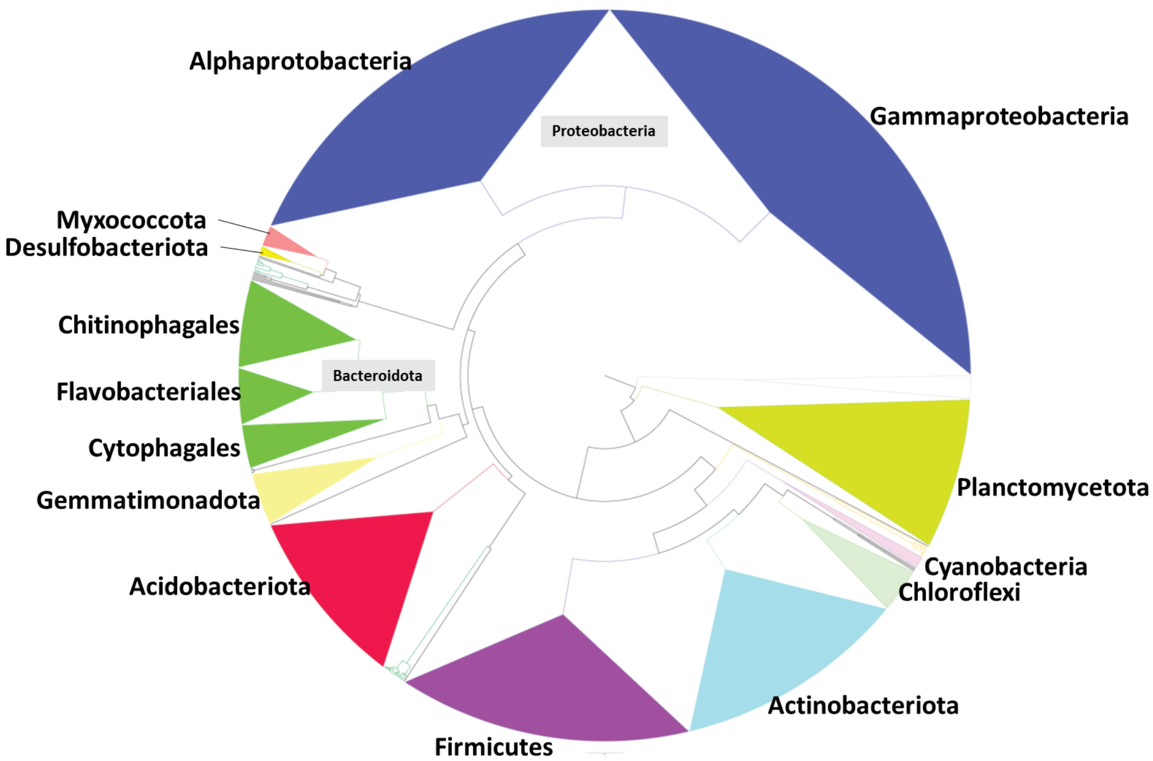


***Appendix 11:*** *Phylogeny of 1,590 ASVs based on full-length 16S rRNA sequencing. Main clades are grouped to simplify comparison with the genome-based phylogeny (Zhu et al., 2019).*
